# *Pseudomonas aeruginosa lasR* mutant fitness in microoxia is supported by an Anr-regulated oxygen-binding hemerythrin

**DOI:** 10.1101/802934

**Authors:** Michelle E. Clay, John H. Hammond, Fangfang Zhong, Xiaolei Chen, Caitlin H. Kowalski, Alexandra J. Lee, Monique S. Porter, Casey S. Greene, Ekaterina V. Pletneva, Deborah A. Hogan

**Author notes:** To whom correspondence should be addressed, Department of Microbiology and Immunology, Geisel School of Medicine at Dartmouth, Rm 208 Vail Building, Hanover, NH 03755. Authors contributed equally to this work (Michelle E. Clay, John H. Hammond, Deborah A. Hogan).

## Abstract

*Pseudomonas aeruginosa* strains with loss-of-function mutations in the transcription factor are frequently encountered in the clinic and the environment. Among the characteristics common to LasR-defective (LasR-) strains is increased activity of the transcription factor Anr, relative to their LasR+ counterparts, in low oxygen conditions. One of the Anr-regulated genes that was highly induced in the LasR-strains encoded a putative oxygen-binding hemerythrin encoded by *PA14_42860* (*PA1673*) which we named *mhr* for microoxic hemerythrin. Purified *P. aeruginosa* Mhr protein contained the predicted di-iron center and binds oxygen with a *K*_d_ of 1 µM. Both Anr and Mhr were necessary for fitness in *lasR*+ and *lasR* mutant strains in colony biofilms grown in microoxic conditions, and the effects were more striking in the *lasR* mutant. Among genes in the Anr regulon, *mhr* was most closely co-regulated with the Anr-controlled high affinity cytochrome *c* oxidase genes and in the absence of high affinity cytochrome *c* oxidase activity, deletion of *mhr* no longer caused a fitness disadvantage suggesting that Mhr works in concert with microoxic respiration. We demonstrate that Anr and Mhr contribute to LasR-strain fitness even in the normoxic biofilm conditions, and metabolomics data indicate that in a *lasR* mutant, expression of Anr-regulated *mhr* leads to differences in metabolism in cells grown on LB and artificial sputum medium. Together these data indicate that increased Anr activity in microoxically-grown *lasR* mutants confers an advantage in part for its regulation of the O_2_ binding protein Mhr.

**Significance:** *Pseudomonas aeruginosa*, a versatile bacterium that both lives in environmental habitats and causes life-threatening opportunistic infections, uses quorum sensing to coordinate gene expression with cell density. The *lasR* gene, which encodes a quorum sensing regulator, is often deleteriously mutated in clinical isolates. Interestingly, LasR-strains have high activity of the oxygen-sensitive transcription factor Anr in microoxic conditions. This report identifies and characterizes an Anr-regulated microoxic hemerythrin that reversibly binds oxygen. We showed both *anr* and *mhr* are critical to fitness in microoxia, and these genes uniquely benefit LasR-strains in normoxia. Our findings enrich our understanding of the success of *P. aeruginosa* as a pulmonary resident through its propensity to lose LasR functionality in the context of low-oxygen infection environments.

## Introduction

*Pseudomonas aeruginosa* is a devastating pathogen for healthcare systems worldwide and causes opportunistic infections at multiple body sites that are extremely difficult to treat. *P. aeruginosa* is especially damaging within the context of the genetic disease cystic fibrosis (CF), where it establishes chronic infections of the airway and is a major predictor of morbidity and mortality (1, 2). The success of *P. aeruginosa* in disease is due to a confluence of factors, including intrinsic and acquired antibiotic resistance (3), production of a battery of secreted, virulence-associated molecules (4), the ability to form antibiotic- and immune cell-recalcitrant biofilms on biotic surfaces and implanted devices (5), and versatile metabolic capabilities (6), such as a pronounced ability to grow in the hypoxic or anoxic conditions engendered by biofilms and chronic infections (7, 8).

*P. aeruginosa* utilizes quorum sensing (QS) to coordinate the expression of a broad set of genes involved in virulence and nutrient acquisition (9). In many different contexts, the gene that encodes one of the key transcriptional regulators involved in QS, *lasR*, frequently sustains loss-of-function mutations. LasR-defective (LasR-) strains of *P. aeruginosa* are commonly isolated from the lungs of individuals with CF (10-12) and other pulmonary diseases (13, 14), implanted device infections (5, 15), acute corneal ulcers (16), and the environment (16).

Several factors may lead to the selection for *P. aeruginosa lasR* mutants and their observed fitness relative to their wild-type counterparts including the advantages of social cheating (17, 18) and enhanced growth on specific carbon and nitrogen sources (9). In addition, both laboratory strains and clinical isolates with mutations in *lasR* display higher expression of the Anr regulon in microoxic environments than their *lasR*-intact counterparts (16, 19). Anr and its homologs have been well-characterized as oxygen-sensitive transcription factors (20-22). The *P. aeruginosa* Anr regulon includes enzymes involved in microoxic and anoxic respiration, fermentation, and CupA fimbriae, as well as biosynthetic enzyme analogs of enzymes that require oxygen and a number of uncharacterized genes including a putative hemerythrin encoded by PA14_42860 (PA1673) (23-25).

Hemerythrins are typified by an antiparallel four-helix bundle with conserved histidine, aspartate and glutamate residues in an H–HxxxE–HxxxH–HxxxxD motif that forms an oxygen-binding di-iron active site (26, 27). Hemerythrins were first characterized as proteins that bind and transport oxygen in the body fluids and tissues of invertebrate worms and leeches (28, 29), and more recent genomic analyses show that hemerythrins are present in all domains of life (27, 30). Many bacterial genomes contain predicted hemerythrins, but roles for these oxygen-binding proteins in bacterial biology have only begun to be characterized (30). A hemerythrin in *Campylobacter jejuni* protects oxygen-sensitive iron-sulfur cluster enzymes from oxidative damage as the cells experience fluctuations in oxygen concentrations (31). In *Methylococcus capsulatus*, a single-domain hemerythrin improves the activity of an inner membrane methane monooxygenase (32), consistent with a role for hemerythrin in oxygen transport and delivery in support of oxic metabolism.

In this work, we demonstrate that in colony biofilms with the wild type, *lasR* mutant fitness is dependent on Anr and the Anr-regulated O_2_ binding hemerythrin Mhr. Mhr is encoded by *PA14_42860* (PA1673), and *mhr* transcript and protein levels are higher upon loss of LasR function in both laboratory strains and clinical isolates. The *P. aeruginosa* Mhr is specifically important for microoxic fitness in the wild type (WT) and confers a fitness advantage in *lasR* mutant colony biofilms under both microoxic and normoxic conditions. The role of Mhr is dependent on the presence of the high affinity cytochrome *c* oxidases suggesting that it supports respiration. Metabolomics data suggest that Mhr contributes to the metabolic differences observed between WT and Δ*lasR* mutants, and its major effects are on central metabolism and amino acid catabolism. These data suggest that microoxic fitness due to Anr-regulated Mhr contributes to the selection for the common, naturally-occurring *lasR* mutants.

## Results

### Anr contributes to Δ*lasR* mutant fitness in microoxic colony biofilms

We previously reported that Δ*lasR* mutants in laboratory strains PA14 and PAO1 as well as in diverse clinical isolates had higher activity of the transcription factor Anr than their LasR+ counterparts under microoxic conditions (16, 19). To determine if Anr activity contributes to the higher fitness of *lasR* mutant strains, we developed an assay for the assessment of fitness in colony biofilms grown under microoxic conditions (0.2%; 2.5 µM O_2_). In competitive fitness assays, wherein different strains were competed with a PA14 strain tagged with a constitutively expressed *lacZ* gene, the two strains had access to a common pool of secreted products including enzymes (e.g. LasA and LasB proteases) and small molecules (such as redox-active phenazines), despite differences in exoproduct production between strains (**Figure 1A** for experimental design). Using this assay, we observed that PA14 Δ*lasR* was more fit than the PA14 wild type under microoxic conditions (**Figure 1B**). The increased fitness of Δ*lasR* could be complemented by restoring *lasR* to its native locus (**Figure 1C**). In both PA14 and the PA14 Δ*lasR* derivative, deletion of *anr* led to a loss of fitness that was complemented by providing *anr* back at the native locus (**Figure 1B**). The presence of *anr* was essential to the microoxic fitness advantage that the Δ*lasR* mutant had over the wild type (**Figure 1B**).

**Figure 1.**
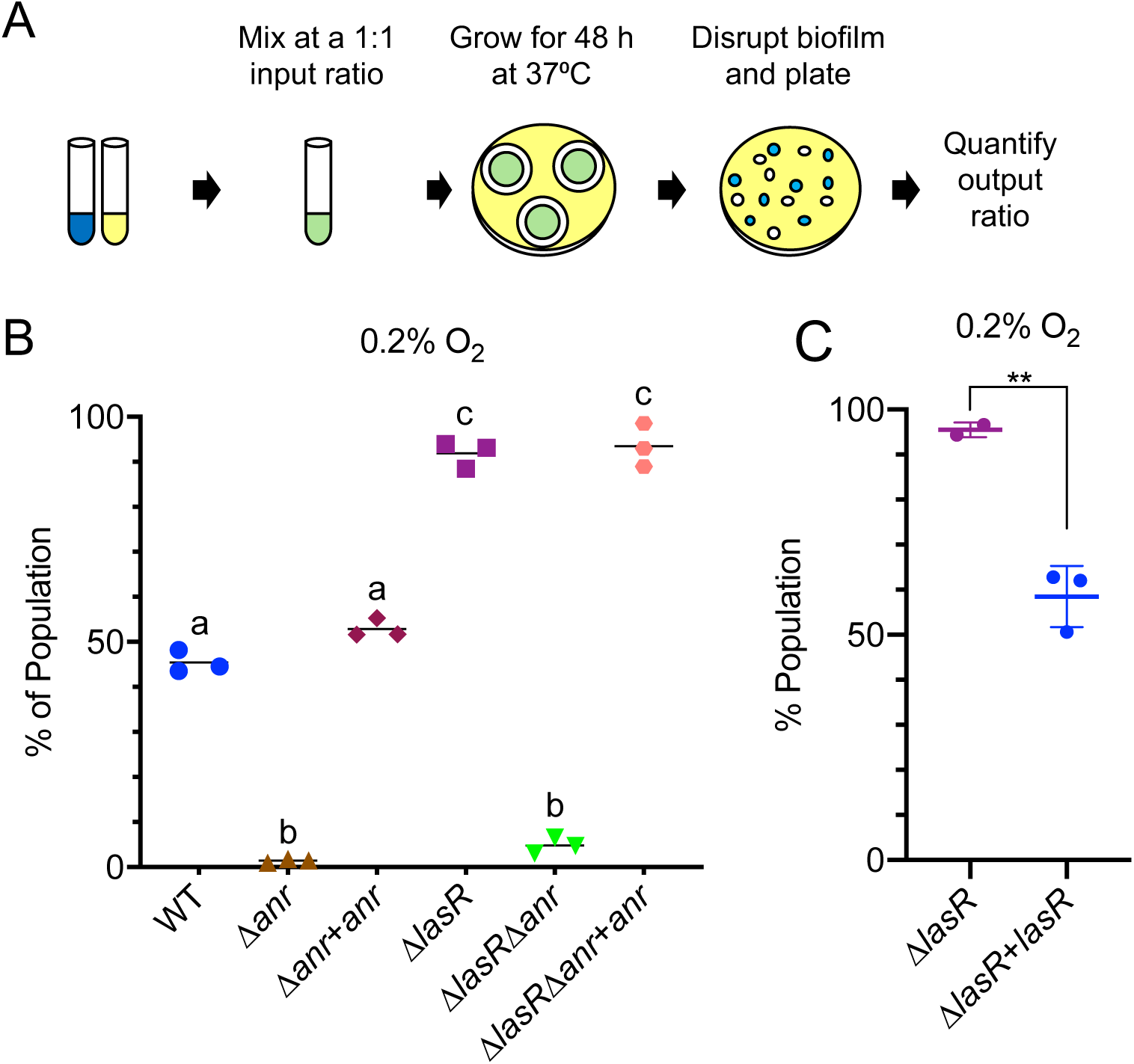
Increased fitness of the Δ*lasR* mutant in microoxic conditions is dependent on *anr*. A) Assay scheme for the competition of *P. aeruginosa* strains (yellow culture) against a *P. aeruginosa* strain tagged with a constitutively-expressed *lacZ* (blue culture). B) Fitness of wild type (WT), Δ*anr*, Δ*anr + anr*, Δ*lasR*, Δ*lasRΔanr*, and Δ*lasR*Δ*anr + anr* relative to the *lacZ*-labeled WT on tryptone agar at 0.2% O_2_ at 37°C. Using a one-way ANOVA with multiple comparisons test, a-b, a-c, and b-c are significantly different, p<0.0001. C) Relative fitness of Δ*lasR* and Δ*lasR* + *lasR* using the assay conditions described for assays in panel B. **, p< 0.01 by unpaired t-test. Relative abundances of strains were determined after 48 h.

### Anr regulates a putative hemerythrin in *P*. *aeruginosa*

The transcription factor Anr regulates dozens of genes involved in denitrification, fermentation and microoxic metabolism. Multiple transcriptome studies comparing *P. aeruginosa* strains to Δ*anr* derivatives in laboratory strain and CF clinical isolate backgrounds revealed that expression levels of the uncharacterized gene *PA14_42860* (*PA1673* in strain PAO1) are severely reduced in the absence of Anr under microoxic and anoxic conditions (7, 19). Furthermore, analysis of pairs of genetically related isolates containing a *lasR* mutant and a *lasR*+ counterpart found this gene, named here *mhr* for <u>m</u>icrooxic <u>h</u>emerythrin, to be one of the most upregulated genes in *lasR* mutants (19). An Anr consensus binding motif (TTGATCGGCGTCAA) was found 81 bp from the translational start of Mhr (33). To test whether Anr was a positive regulator of *mhr*, we constructed a *mhr* promoter fusion to *lacZ* and integrated it at a neutral site on the chromosome in the WT and Δ*anr* backgrounds. Under microoxic conditions, *mhr* promoter activity was 64-fold higher in the wild type than in the Δ*anr* mutant. Mutation of two nucleotides in the Anr consensus motif that are essential for Anr regulation of other promoters (34) led to a large and significant reduction in β-galactosidase production (**Figures 2A**). Consistent with previous reports that *mhr* transcripts were more abundant in *lasR* loss-of-function mutants compared to their *lasR*-intact counterparts, *mhr* promoter activity was 1.9-fold higher in the Δ*lasR* mutant compared to the wild type (**Figure 2B**). As in the wild type, promoter activity was Anr-dependent in the Δ*lasR* mutant (**Figure 2B**).

**Figure 2.**
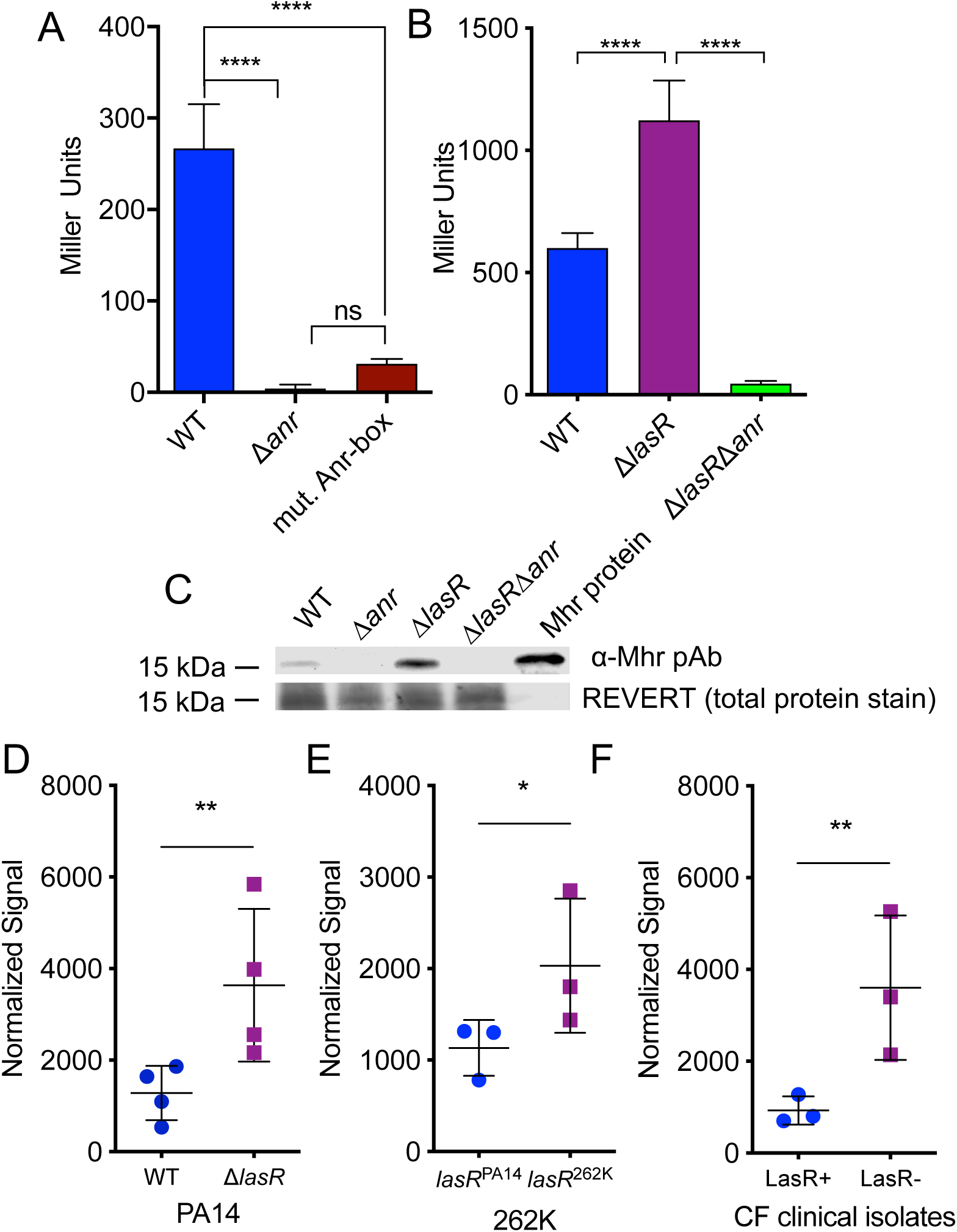
*mhr* transcription and Mhr protein levels are Anr-regulated and higher in the *lasR* mutant strains. **A-B)** *β*-galactosidase activity in strains bearing an *mhr-lacZ* promoter fusion. A) PA14 wild-type (WT), Δ*anr*, and WT bearing a promoter fusion variant in which the Anr-box mutated as described in the methods (mut. Anr-box). B) *β*-galactosidase activity in WT, Δ*lasR*, and Δ*lasR*Δ*anr* containing the *mhr-lacZ* promoter fusion. ****, p<0.0001 by oneway ANOVA, multiple comparisons test; ns, not significant. C) Western blot using a rabbit α-Mhr polyclonal antibody. Lanes from left to right: WT, Δ*anr*, Δ*lasR*, Δ*lasR*Δ*anr* and 12.5 ng Mhr that was purified from *E. coli*. Total protein stained with REVERT, used for normalization, is shown. D-F) Quantification of Mhr by Western blot for pairs of *lasR* mutant and *lasR*+ strains, normalized to total protein using the REVERT stain (Licor). Each point within a group represents band intensity data from a separate experiment on a different day. *p<0.05, **p<0.01 by ratio paired t-test. D) Mhr levels in PA14 WT and Δ*lasR*. E) Mhr levels in the natively LasR-keratitis isolate 262K and a derivative in which its *lasR* allele was replaced with that from strain PA14. F) Mhr levels for a LasR+ chronic CF isolate and its geneticallyrelated LasR-partner recovered from the same patient. Cells were grown as colony biofilms in 0.2% oxygen for 16h on tryptone agar for all experiments in this figure.

To determine if Mhr protein levels were also higher in LasR-strains, we expressed and purified *P. aeruginosa* Mhr from *E. coli* (**Figure S1**) and used it to raise an anti-Mhr polyclonal antibody. Western blot analysis detected purified Mhr at about 15 kDa, similar to its predicted molecular weight of 17.9 kDa, and a similar band was detected in *P. aeruginosa* whole cell lysates from the wild type but not in the Δ*anr* mutants (**Figure 2C**). Mhr levels in the Δ*lasR* strain were also dependent on *anr* at the protein levels. We quantified the relative amounts of Mhr using the Licor Odyssey near-IR Western blotting system. Mhr protein levels were 2.8-fold higher in the Δ*lasR* mutant than in the wild type (**Figures 2D**). We also analyzed Mhr levels in a clinical isolate from a corneal eye infection (262K) that was a natural *lasR* mutant (16) and a 262K derivative in which the native allele was replaced with the PA14 allele, as well is two clinical isolates from the same subject with CF with one isolate bearing a *lasR* mutation. We found that in the two clinical isolate backgrounds, the *lasR* mutant in each pair had significantly more Mhr than the corresponding LasR+ strain (**Figures 2E-F**).

### Mhr is a bacteriohemerythrin that reversibly binds oxygen via a di-iron center

The Mhr sequence contains the conserved di-iron binding motif H–HxxxE–HxxxH–HxxxxD (26, 27). When the *P. aeruginosa* Mhr sequence was threaded onto the crystal structure of *Methylococcus capsulatus* hemerythrin (Hr), all seven of the residues were predicted to form the metal-binding active site (**Figure 3A**). In addition to its presence in other *Pseudomonas* spp., the closest Mhr homologs by sequence were found in other gamma proteobacteria (30) (*Stenotrophomonas maltophilia, Xanthomonas campestris, Acinetobacter baumannii, Dyella* spp.) and the nitrogen-fixing plant symbiont *Azobacter chroococcum*, and the active site residues are conserved across all of these homologous proteins (35) (**Figure S2**). Consistent with presence of the di-iron binding motif, we detected iron in Mhr protein at a ratio of 2:1 (see methods for description of the assay).

**Figure 3.**
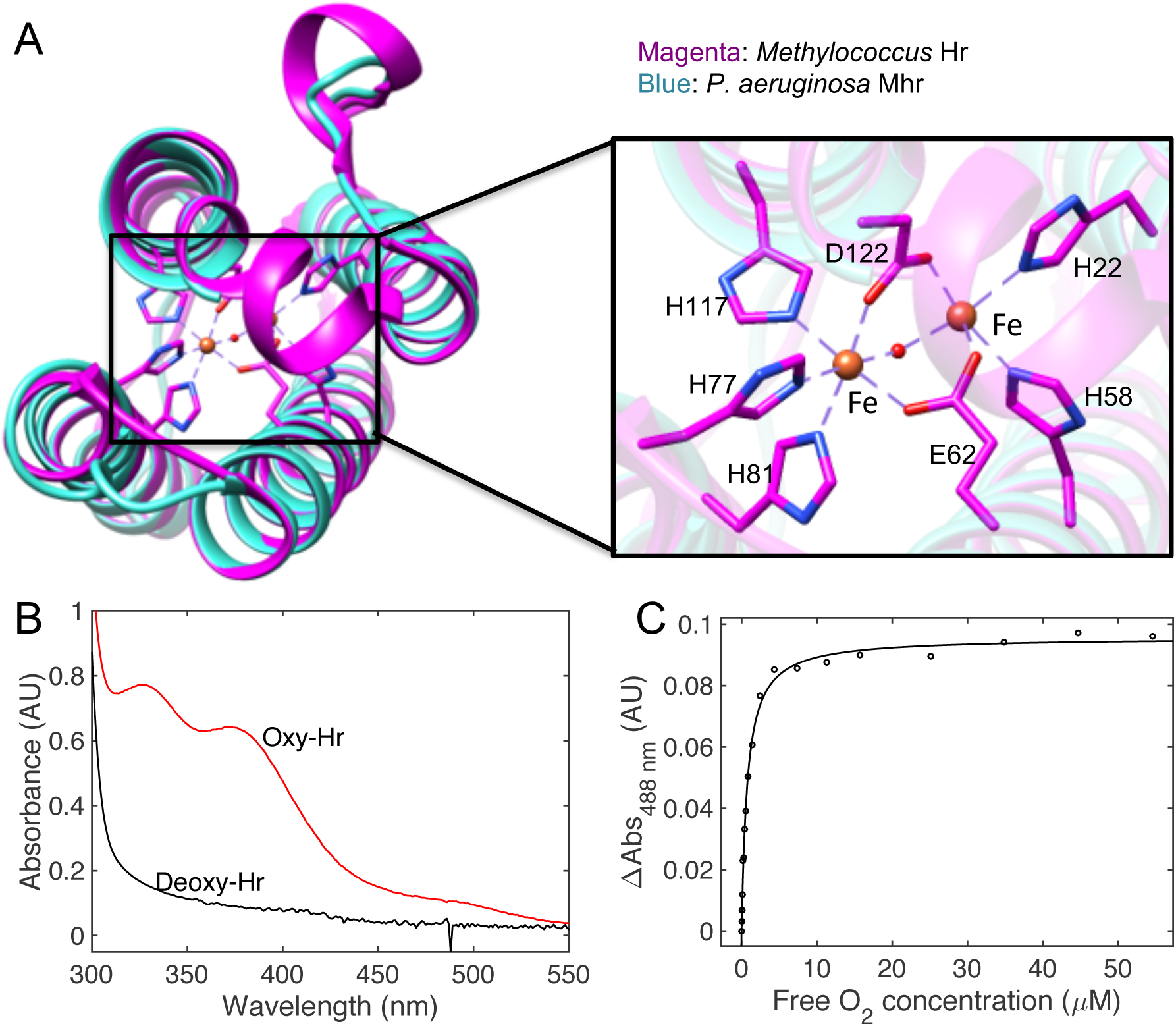
Mhr is an oxygen-binding hemerythrin. A) Structural model of *P. aeruginosa* Mhr based on the crystal structure for *Methylococcus capsulatus* hemerythrin protein (*Methylococcus* Hr) (4xpx.pdb). The region around the di-iron center was zoomed in to display the key oxygen binding residues. B) Electronic absorption spectra of deoxy-Mhr (Fe^2+^Fe^2+^) and oxy-Mhr (Fe^3+^Fe^3+^-O_2_) in 20 mM Tris-HCl, pH 8. The broad peak around 488 nm is characteristic of the oxy-Mhr form. C) The plot of the absorbance change with the free O_2_ concentration was fitted to indicate the apparent O_2_ binding constant (*K*_d_ = 0.74 μM).

Mhr purified from *E. coli* had an absorbance spectrum with maxima similar to *M. capsulatus* Hr (peaks at 328nm and 371nm) and other hemerythrin proteins (36). As in other hemerythrins, there were spectroscopic differences between the oxygen bound (oxy-Mhr) and oxygen-free species (deoxy-Mhr) (**Figures 3B and S3A**). Analysis of spectroscopic changes upon titration of oxygen showed an apparent *K*_D_ for O_2_ of 0.74 µM (**Figures 3C and S3B**). Thus, Mhr was shown to bind iron and oxygen with an affinity that is relevant to microoxic conditions.

### Mhr plays a role in fitness in microoxic conditions but not anoxic conditions

To test whether Mhr was important for growth under microoxic conditions, we constructed Δ*lasR*Δ*mhr* and Δ*mhr* strains and performed competition assays at 0.2% oxygen. Both the Δ*mhr* and the Δ*lasR*Δ*mhr* mutant were less fit than their parental strains under microoxic conditions (**Figure 4A**). We were able to complement the fitness phenotypes of the Δ*mhr* strains by expressing *mhr* using the arabinose-inducible plasmid pMQ70 (**Figure 4B-C**). Under anoxic conditions in which nitrate is used an alternative electron acceptor the Δ*mhr* had no fitness defect, suggesting that Mhr, an oxygen binding protein, does not play a physiological role in the absence of oxygen (**Figure S4**).

**Figure 4.**
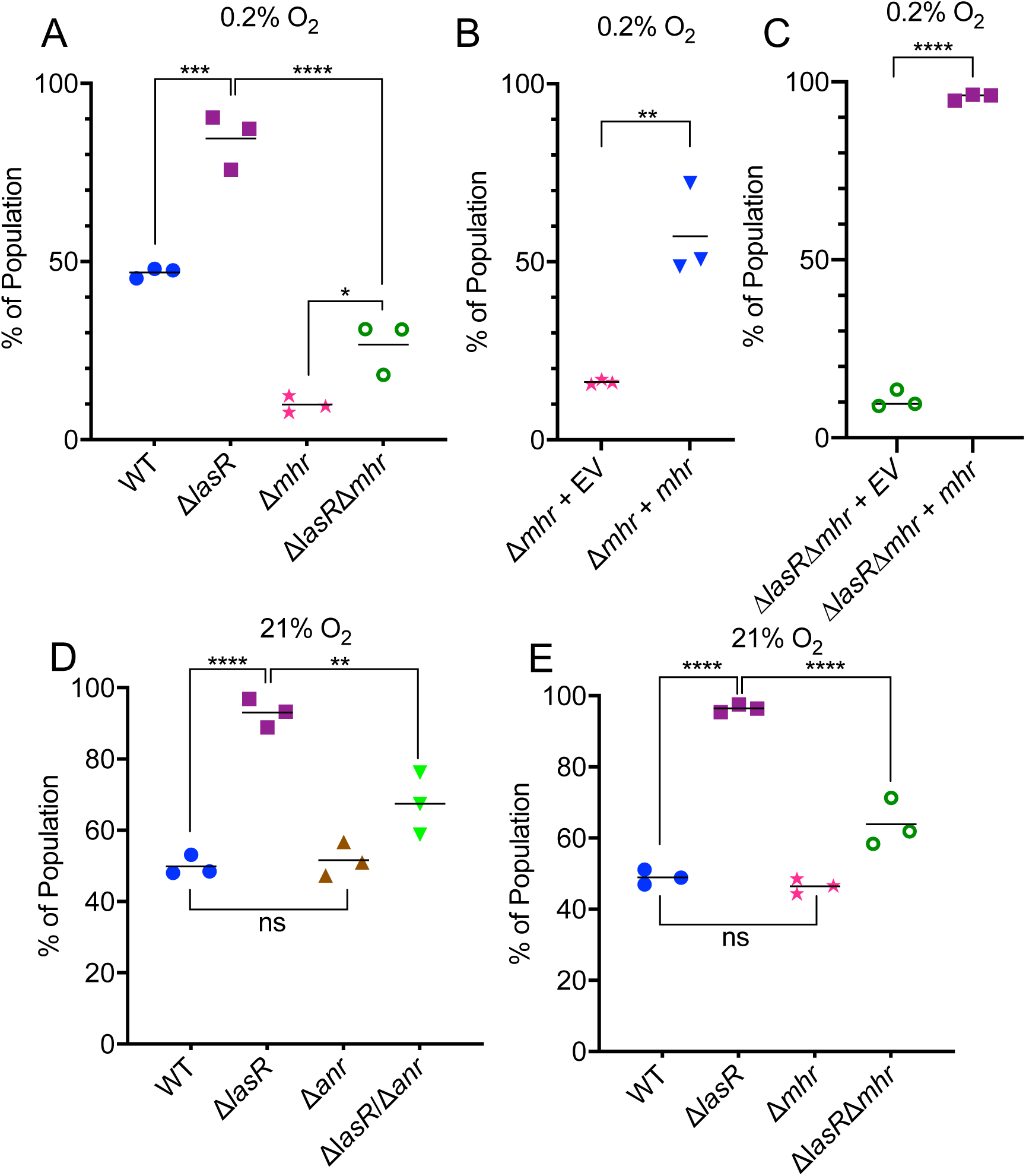
Fitness of the Δ*lasR* mutant in microoxic and normoxic conditions is dependent on *mhr*. A) Microoxic fitness of the WT, Δ*lasR*, Δ*mhr*, and Δ*lasR*Δ*mhr* relative to the *lacZ*-labeled WT under 0.2% O_2_. B) Microoxic fitness of Δ*mhr* carrying the empty vector pMQ70 (EV) or a pMQ70-mhr (*mhr*) against the *lacZ*-labeled WT carrying pMQ70. C) Microoxic fitness of the Δ*lasR*Δ*mhr* + EV or Δ*lasR*Δ*mhr* + *mhr*. In B and C, 0.2% arabinose and carbenicillin (300μg/ml) was added to the agar medium. D) Normoxic (21% O_2_) fitness of WT, *ΔlasR, Δanr*, and Δ*lasR*Δ*anr*. E) Normoxic fitness of *ΔlasR, Δmhr*, and Δ*lasR*Δ*mhr*. All assays were performed grown on tryptone agar at 37°C for 48 h. (*, p≤0.05, ***, p≤0.001, ****, p≤0.00001 by one-way ANOVA, multiple comparisons (A, D and E) or t-test (B and C).

### Mhr plays a role in Δ*lasR* mutant fitness in normoxic conditions

Under normoxic conditions (21% oxygen), as in microoxic conditions, the Δ*lasR* mutant was more fit than the wild type in our colony biofilm competition assay (**Figure 4D**). Deleting *anr* in the wild type did not affect fitness under normoxia, but it did result in a fitness defect in the Δ*lasR* background (**Figure 4D**). Similarly, the Δ*mhr* mutant equally was as fit as the wild type under normoxic conditions (**Figure 4E**), but loss of *mhr* led to reduced fitness in the Δ*lasR* mutant background. Together, these data suggest that Anr, in part due to its regulation of Mhr, contributes to Δ*lasR* fitness even in colony biofilms grown in normoxia.

The role for Mhr in PA14 Δ*lasR* but not wild type in normoxic colonies raised the question of whether Anr activity is more strongly induced in Δ*lasR* mutants as oxygen becomes depleted. Using a liquid culture assay in which increasing volumes of medium were used to decrease the surface area for oxygen transfer, we found that the *mhr* promoter induction was low in 1 ml cultures and it increased with culture volume which is consistent with Anr activity increasing as oxygen availability decreases (**Figure S5A**). At each volume, the Δ*lasR* mutant had significantly higher levels of *mhr* promoter activity than the wild type while culture densities were similar (**Figure S5B**). Using the 10 ml volume, we found that like Δ*lasR*, the Δ*lasI* strain which cannot synthesize the LasR ligand 3OC12HSL also had significantly higher levels of *mhr* promoter activity and the elevated activity could be complemented by exogenous 3OC12HSL (**Figure S5C**).

### *mhr* is co-regulated with and epistatic to genes involved in microoxic respiration

To gain more insight into the role of Mhr in microoxic fitness and in light of the finding that the Δ*mhr* mutant does not have a fitness defect in anoxic denitrifying conditions (**Figure S4**), we analyzed its expression pattern relative to Anr-regulated genes known to be involved in metabolism. To do so, we used eADAGE, a tool that enables the visualization of transcriptional pattern relationships deduced from the analysis of 1056 publicly available samples(37). Using the Anr-regulated genes shown in color in **Figure 5A** as input, we found that *mhr* showed an expression pattern that closely mirrored those of genes that encode components of the CcoNOPQ-2 high affinity cytochrome *c* oxidase. Thus, we sought to assess the genetic interaction between Mhr and the high affinity cytochrome *c* oxidases. The loss of the *cbb*_3_-type high affinity cytochrome *c* oxidase encoded by *ccoNOPQ-2* has little effect on microoxic growth (7) because of the presence of *ccoNOPQ-1* which encodes a highly similar enzyme. A strain lacking both *cco* operons (Δ*cco*) has a significant microoxic growth defect (7) suggesting that the two *cbb*_3_-type oxidases have partially redundant or compensatory activities. To determine if the Mhr function required high affinity cytochrome *c* oxidases, we competed the *lacZ*-tagged Δ*cco* strain against either an untagged version of itself or the untagged *ΔccoΔmhr* mutant. The Δ*cco* and Δ*cco*Δ*mhr* mutants had similar fitness profiles, suggesting that Mhr either works in concert with the high affinity cytochrome *c* oxidases or that these oxidases are important for creating the environment in which Mhr is most important (p=0.1963) (**Figure 5B**).

**Figure 5.**
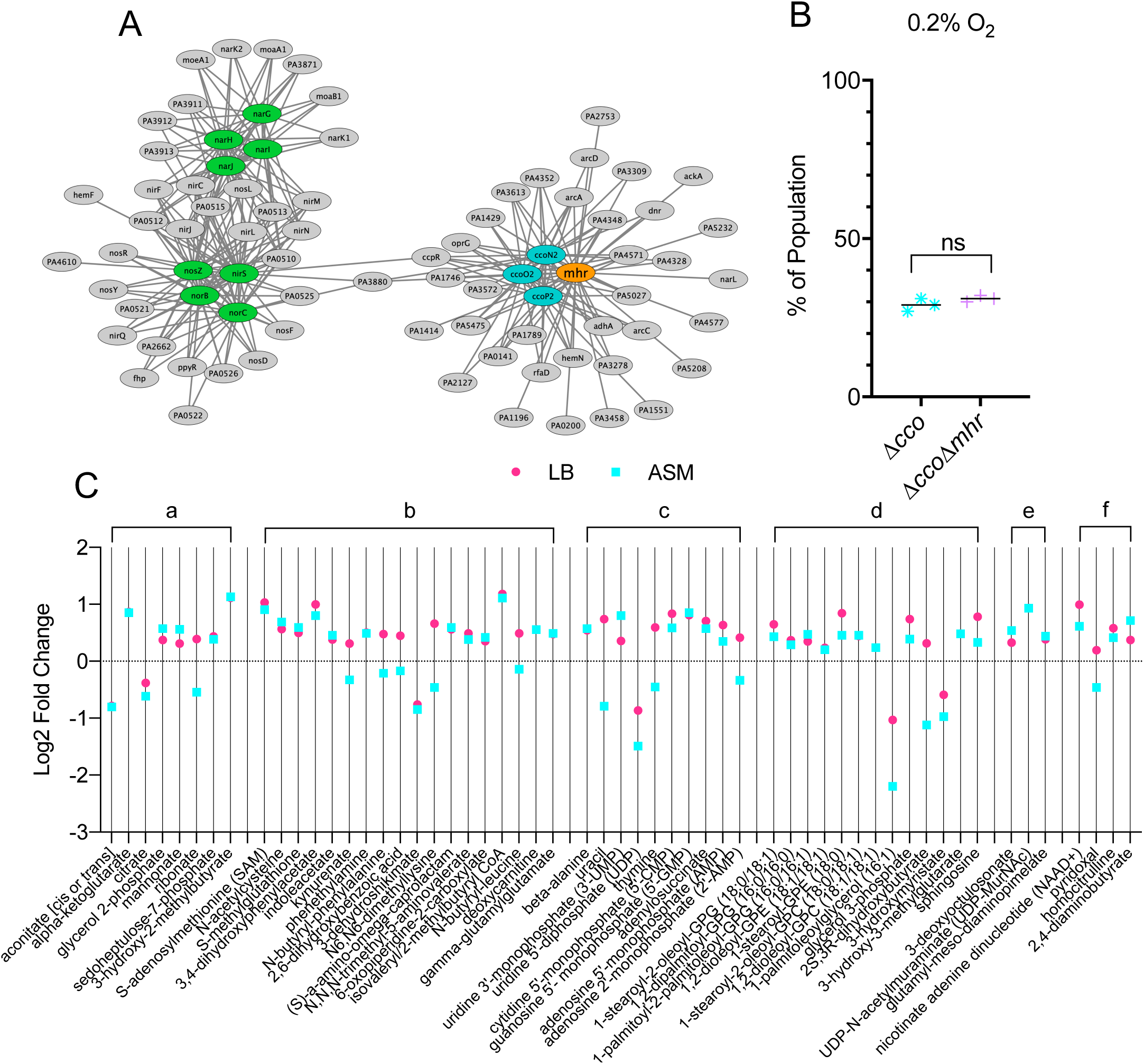
*mhr* is co-regulated with and epistatic to the *ccoNOPQ-2* high affinity cytochrome *c* oxidase and affects cell metabolism. A) Similarity in expression pattern for genes within the Anr regulon in a publicly available data compendium determined using a machine learning derived model for gene expression, eADAGE. Similarities of expression pattern for Anr-regulated genes (ovals) are depicted as edge length. High affinity cytochrome C oxidases, *ccoN2, ccoO2, ccoP2*, are shown in blue and denitrification structural genes are shown in green. B) Microoxic (0.2% O_2_) fitness comparison for Δ*cco*1Δ*cco*2 and Δ*cco*1Δ*cco*2Δ*mhr* was determined by competition against the constitutively *lacZ*-labeled Δ*cco*1Δ*cco*2. Assay was performed on tryptone agar at 37°C for 4 d. C) Metabolomics analysis of 16 h colony biofilms on either LB agar or artificial sputum medium (ASM) agar under 21% oxygen. Log_2_ transformed fold changes (Δ*lasR*/Δ*lasR*Δ*mhr*) were plotted if they met the following criteria: 1) p<0.1 by Welch’s t-test (unequal variance) and 2) metabolite was detected on both media. Plotted metabolites were grouped by pathway: a) energy generation, b) amino acid metabolism, c) nucleotide metabolism, d) lipid metabolism, e) LPS and cell wall components, f) cofactors, vitamins, and urea cycle. Each point represents the average of five replicates.

### Metabolomics analysis indicates that Mhr influences the metabolism of a Δ*lasR* mutant

In light of the contribution of Mhr to the fitness of Δ*lasR* under normoxic conditions, we assessed the effects of Mhr on the metabolism of the Δ*lasR* mutant by performing a metabolomics comparison of Δ*lasR* and Δ*lasR*Δ*mhr* colony biofilms grown with atmospheric oxygen. We analyzed cells grown on LB and on artificial sputum medium designed to mimic the sputum conditions found in the lungs of individuals with CF, referred to as artificial sputum medium(38) and focused on compounds that were significantly different upon the absence of Mhr on both media types. Our analysis found that comparison of Δ*lasR* to the Δ*lasR*Δ*mhr* mutant across both media revealed significant differences in fifty-seven metabolites (**Figure 5C** and **Table S1**). Among the largest differences between Δ*lasR* and Δ*lasRΔmhr* were those metabolites involved in energy generation, including metabolites in the TCA cycle. Amino acid metabolic pathways, particularly those involved in the metabolism of sulfur-containing and aromatic amino acids, were also affected by Mhr. Other pathways included those related to the metabolism of nucleotides, lipids, LPS and cell wall, cofactors and vitamins. The differences detected in these pathways are consistent with Mhr affecting metabolism and oxic respiration in the cell.

## DISCUSSION

Findings from this study and others demonstrate that *P. aeruginosa* LasR-strains have high Anr activity (16, 19) and activity of Anr-regulated pathways (39) in microoxic environments. In this study, we showed that Δ*lasR* mutants had increased fitness in the colony biofilms grown in microoxic and normoxic atmospheres, and that this increase in fitness was dependent on *anr* and Anr-regulated *mhr*. Furthermore, the relative fitness defect incurred by loss of either *anr* or *mhr* was greater in a Δ*lasR* mutant background than in the wild type (LasR+). Compared to other Anr regulated genes, expression of *mhr* and the Anr-regulated *ccoNOPQ-2* high affinity terminal oxidase followed similar patterns across a large data compendium and the loss of Mhr did not cause a further fitness defect in a strain lacking high affinity cytochrome *c* oxidase activity, suggesting that Mhr and high affinity cytochrome *c* oxidases work together directly or indirectly. Both proteomic and transcriptomic studies provided evidence for the coordinated induction of Mhr and high affinity cytochrome *c* oxidases (40) in synthetic cystic fibrosis medium under microoxic conditions compared to normoxic conditions, and in CF sputum (40, 41).

There are several possible models for how Mhr improves microoxic fitness. Hemerythrins have been shown to play a role in oxygen transport and delivery (42) and our observation that *mhr* is tightly co-expressed with the *cbb*_3_-2 high affinity cytochrome *c* oxidase (*ccoNOPQ*-2) suggests that Mhr could play some mechanistic role in improving access of the cytochrome *c* terminal oxidases to oxygen beyond what would be achievable via passive diffusion when extracellular oxygen levels are low. In *Methylococcus capsulatus (32)*, a homologous hemerythin delivers O_2_ to methane monooxygenase, which resides on intracytoplasmic membrane. Future studies will determine if the C-terminal domain that is conserved in *Pseudomonas* Mhr homologs (and absent in other types of single-domain hemerythrins) plays a role in protein-protein or Mhr-membrane interactions in ways that promote microoxic respiration by high affinity *cbb*_3_ cytochrome *c* oxidases.

Our metabolomics data suggest that Mhr influences the metabolic state of the cell, perhaps through direct binding of oxygen and altering O_2_ availability to metabolic oxygenases or conferring protection to O_2_ sensitive metabolic enzymes. The perturbation of metabolites in the TCA cycle upon mutation of *mhr* may be consistent with a role in central metabolism; there was also a striking effect on amino acid catabolism and in particular, enzymes involved in sulfur metabolism which may be related to changes in intracellular redox. This finding suggests that Mhr could perhaps function to stabilize oxygen-sensitive proteins that are active in microoxic conditions, similar to the hemerythrins of the microaerophilic *Campylobacter jejuni* that protect iron-sulfur cluster proteins during exposure to high oxygen concentrations (31).

The elevated Anr activity in *lasR* mutants was not abolished in co-culture by the presence of factors produced by the wild type. The model that Δ*lasR* mutants have a fitness advantage due to elevated Anr activity, and therefore high Mhr, is not incompatible with models for increased Δ*lasR* mutant fitness relative to *lasR*+ strains due to the benefits of social cheating and the cost-less exploitation of public goods (17). Other studies have shown LasR-strains have increased resistance to lysis in alkaline conditions (43) and enhanced growth on particular carbon sources(9) which likely also promote fitness. Thus, we posit that oxygen limitation can act as one positive selection pressure against LasR signaling. We do not fully understand how an increase in Anr-activity is incurred by *lasR* deficiency, and further studies are needed to understand the molecular mechanism behind this phenomenon.

## MATERIALS AND METHODS

### Bacterial strains and growth conditions

All bacterial strains and plasmids used in this study are listed in **Table S2**. Bacteria were routinely grown in lysogeny broth (LB) at 37°C, except in the case of *E. coli* transformed with pUT18, which was grown at 30°C, or during protein purification, as described below. For microoxic studies, bacteria were grown inside a hypoxic cabinet with O_2_ and CO_2_ controllers (COY Laboratory Products, Grass Lake, MI), at 0.2% O_2_ and 5% CO_2_. For experiments done in anoxic conditions bacteria were grown in an air-tight chamber, and oxygen was depleted using a GasPak catalyst and indicator system (BD 260001). Tryptone broth (TB; 1% tryptone and 0.5% NaCl) and tryptone agar (TA; TB amended with 1.5% agar) were used for experiments where noted.

### Plasmid construction

Plasmids for making in-frame deletions, complementation, for gene expression, and for integration of the *mhr* promoter fusions into *P. aeruginosa* were constructed using a *Saccharomyces cerevisiae* recombination technique described previously using primers listed in **Table S2**(44). Plasmid pET20b(+)_*mhr* for expression and purification of Mhr was constructed by amplifying *mhr* from template plasmid pMQ70_*mhr*_exp. The PCR product was purified via gel extraction, digested with *Nde*I and *Xho*I, and ligated into the pET20b(+) vector (*Nde*I, *Xho*I). The ligation product was transformed into *E. coli* XL1Blue super-competent cells (Agilent) for propagation. All plasmids were sequenced at the Molecular Biology Core at the Geisel School of Medicine at Dartmouth.

### Biofilm competition experiments

Competition assays were performed to determine relative fitness of *P. aeruginosa* mutants. Strains to be competed were grown overnight in antibiotic selection when appropriate. 1 mL culture was pelleted, resuspended in TB and adjusted to OD_600_ = 0.5. Strains to be competed were combined in a 1:1 ratio and 5 µL of the combined suspension was spotted onto a 0.22 µM polycarbonate filter (Millipore) placed on the surface of a TA plate (supplemented with 0.2% arabinose and/or 300 μg/ml carbenicillin when appropriate) and incubated at 37°C. Filters were transferred to fresh plates after 24 h, and after 48 h of total incubation time, each filter was lifted from the plate and transferred into a 1.5 mL tube. The filter-associated biofilm was disrupted by adding 1 ml PBS + 0.01% Triton X-100 detergent and agitating the tubes at high speed for 2 minutes using a Genie Disruptor (Zymo). This suspension was diluted, spread on LB plates supplemented with 150 µg/mL 5-bromo-4-chloro-3-indolyl-β-D-galactopyranoside (X-Gal) using glass beads, and incubated at 30°C until blue colonies were visible (∼24h), and the number of blue and white colonies per plate were counted and recorded.

### Promoter fusion experiments

For the promoter fusion experiments, the cells were grown in 5 ml LB overnight as described above. For colony biofilm-based assays, overnight cells were washed, normalized to OD_600_ = 0.5, and spotted onto 0.22 µM polycarbonate filters (Millipore) on TA. These cells were incubated for 16 h in the microoxia chamber (0.2% O_2_) at 37°C and then harvested into 1 ml of PBS. For flask-based assays, overnight cells were adjusted to OD = 0.05 in TB and the noted volume (20, 10, 5, 1 ml) was distributed into a sterile 50 ml capacity Erlenmeyer flask covered loosely with foil. These flasks were incubated at 37°C while shaking at 225 rpm for 6 h. Harvested cells were evaluated for β-galactosidase production as described by Miller(45). Briefly: Harvested cells were placed on ice for 20 minutes, then 100 μl of cells were distributed in triplicate into wells of a 96-well plate for measuring and recording OD_600_ and in triplicate into 13×100mm glass tubes for measuring β-galactosidase activity. Z-buffer (60 mM Na_2_HPO_4_, 40 mM NaH_2_PO_4_, 10 mM KCl, 1mM MgSO_4_, 50 mM β-mercaptoethanol, pH 7.0) was added (900 μl) to the glass tubes, followed by 50 μl chloroform and 25 μl SDS (0.1%) to kill and perforate the cells. The tubes were vortexed and then incubated at 30°C for 5 minutes, followed by addition of 200 µl of a Z-buffer based stock solution of o-Nitrophenyl-β-D-galactopyranoside (ONPG, 4mg/ml). After yellow color developed, 500 µl of stop solution (1 M Na_2_CO_3_) was added to each tube, and the time elapsed since ONPG addition was recorded. Aliquots (100 μl) of this solution were transferred to a 96-well plate and the absorbance at 420nm and 550nm was recorded. Miller units were calculated using the equation: Miller Units = 1000 × [(OD_420_ – (1.75 × OD_550_)) / (T × V × OD_600_)], where T is time elapsed in minutes and V is volume of culture added (100 μl).

### Protein expression and purification

Protein expression and purification techniques were based on protocol by Miner et al. (46). Briefly, pET20b(+)_*mhr* was transformed into *E. coli* BL21(DE3) (Agilent) for expression and grown overnight at 37°C in 30 ml LB supplemented with 100 μg/ml ampicillin. *E. coli* cells were then sub-cultured into six 1 L flasks containing LB, and inoculated flasks were cultured at 37°C with shaking (220 rpm) until OD_600_ = ∼0.8 (roughly 6 hours). The temperature was then lowered to 25°C and 0.2 mM isopropyl-β-D-thiogalactopyranoside (IPTG) was added to induce *mhr* expression. Ferrous ammonium sulfate (100 mg/L) was also added to the media at this time to ensure that sufficient iron was available for synthesis of the Mhr holoprotein. This culture was incubated at 25°C with shaking for 20 h, then the cells were harvested by centrifugation at 5000×*g* for 25 minutes. Cells were re-suspended in 150 ml of HisTrap binding buffer (25 mM Tris-HCl pH 7.8, 500 mM NaCl, 25 mM imidazole, 10% glycerol) and lysed by French Press. The supernatant was separated from cell debris by centrifugation at 22,000×*g* for 45 minutes. Supernatant was loaded onto a 5 ml HisTrap column (GE Healthcare) pre-loaded with Ni^2+^, and Mhr was eluted by HisTrap eluting buffer (25 mM Tris-HCl pH 7.8, 500 mM NaCl, 500 mM imidazole, 10% glycerol). Purified protein was immediately exchanged into TEV cleavage buffer (50 mM Tris-HCl pH 8.0, 10% glycerol) using a 50 ml desalting column (GE Healthcare). For His-tag cleavage, 2 mM DTT, 0.5 mM EDTA and TEV (Mhr:TEV = 10:1) were added. The cleavage reaction was performed at room temperature for 2 h, and then the protein was exchanged into HisTrap buffer and purified by HisTrap column (column flow-through was collected this time). Immediately prior to biochemical experiments using purified Mhr, protein was further purified using a 5 ml Q column (GE Healthcare). Protein was exchanged into 10 mM Tris-HCl pH 8.0 buffer, loaded onto Q column, and eluted by gradually increasing the ionic strength of the buffer (0 to 1M NaCl in 180 minutes). Fractions containing hemerythrin protein were collected, concentrated, and used in subsequent assays.

### Western Blot for Mhr levels

Strains were grown as colony biofilms on tryptone agar at 37°C for 16 h at 0.2% (2.7μM) oxygen. Cells were scraped from the surface of the agar using a glass speader and harvested into 2 ml of PBS. The cells were adjusted to a final OD_600_ of 10 in Laemmli sample buffer (4% SDS, 20% glycerol, 120 mM Tris HCl (pH 6.8) prepared as a stock solution, then 0.02% bromophenol blue and 2-5% β-mercaptoethanol spiked into freshly prepped samples), because we were unable to precisely normalize the total protein between WT and Δ*lasR* background strains using a BCA assay. Samples were heated at 95°C for 10 minutes prior to loading into gels. Five to ten microliters of BioRad All Blue Precision Plus molecular weight marker, purified Mhr, and cell lysate samples were loaded into a 4-15% BioRad pre-cast gradient gel. The gel was run for 10 min at 60 V, followed by ∼40 minutes at 150 V. The BioRad Turboblot semi-dry transfer system (low MW preset) was used to transfer proteins to a LF-PVDF membrane (Immobilon Product IPFL00010). The blots were processed using the standard LICOR protocol for Western blotting, including the optional drying step after transfer and REVERT total protein staining. Our anti-Mhr polyclonal antibody, custom raised in rabbits by Cocalico Biologicals (Stevens, PA) was diluted 1:5,000. The goat anti-rabbit secondary antibody labeled with IRDye 800CW (Licor) was diluted 1:15,000. Blots were imaged using an Odyssey CLX scanner (Licor) and analyzed using the Empiria software (Licor).

### Generating absorption spectra for met-, deoxy-, and oxy-Mhr

Isolated *P. aeruginosa* Mhr was oxidized by at least a 10 molar equivalent of potassium ferricyanide at 4°C for 16 h. Excess oxidant was removed by a PD-10 desalting column (GE Healthcare) equilibrated with 20 mM HEPES, pH7.4 for the resulting met-Mhr. Isolated Mhr (∼200 μM) was reduced to deoxy-Mhr by sodium dithionite (20 molar equivalents) and 1 mM methylviologen (5 molar equivalent) under a nitrogen atmosphere in a nitrogen-filled glove box (COY Laboratory Products) for 3 h. Excess reducing agents were removed by running the protein solution through a PD-10 column (GE Healthcare) equilibrated with 20 mM HEPES, pH 7.4 in the glove box. Oxy-Mhr was captured by exposing deoxy-Mhr to air for 1 min. Absorption spectra were acquired on an Agilent 8453 diode-array spectrophotometer at room temperature.

### Characterizing oxygen binding by Mhr

Deoxy-Mhr protein solution (50-100 μM) was transferred into a cuvette and sealed by a septum cap in the glove box. The cuvette carrying the sample was taken out of the glove box for the absorption spectral measurements using an Agilent 8453 spectrophotometer. Small aliquots of air-saturated buffer (20 mM HEPES, pH 7.4) were injected with a syringe into the protein solution through the septum cap stepwise and the absorption spectra were collected immediately following each addition of the buffer. Absorbance changes at 488 nm were plotted against the oxygen concentration and the data were fit to the following equation from Wendt et al. (47) using the MATLAB curve-fitting tool (MathWorks):

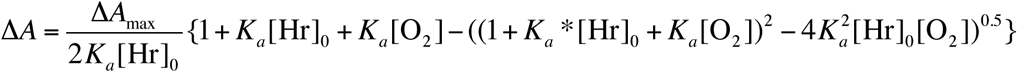

### Sequence alignment of Mhr homologs

Amino acid sequence alignment was performed using ClustalW (35). Sequence identity percentage was determined using Mhr as the reference sequence.

### Iron content determination

The iron content was obtained using a standard ferrozine assay (47). The protein concentration was determined by Bradford protein assay using bovine serum albumin as the standard.

### Structural Homology Model of Mhr

The *P. aeruginosa* Mhr structural homology model was created by the SWISS-MODEL web server (http://swissmodel.expasy.org/) using the structure of *M. capsulatus* hemerythrin (Protein Database ID: 4XPX) as the template. The superimposition of both structures was generated in Chimera (48).

### Using eADAGE to visualize a compendium-wide profile of Anr-regulated gene expression patterns

Selected Anr-regulated genes (*narGHIJ, norBC, nirS, nosZ, ccoNOP*-2, and *mhr*) were applied as input using the “Ensemble ADAGE 300 with more complex gene-gene network” web tool (37) (https://adage.greenelab.com). The “Explore genes’ model similarities” option was used and the genes were viewed as a network. Gene edge weights were downloaded used in conjunction with Cytoscape (49) to color the gene network for clarity.

### Metabolomics

Strains were grown as colony biofilms on LB agar or artificial sputum medium (ASM) agar modified from Palmer, et al (38). In brief, 2X buffered base containing 2.5mM NaH_2_PO_4_ monohydrate (stock 0.2M), 2.5mM Na_2_HPO_4_ (stock 0.2M), 0.7mM KNO_3_ (stock 1M), 0.54mM K_2_SO_4_ (stock 0.25M), 8mg/ml yeast synthetic dropout -Trp (Sigma Y1876), 6.06mg/ml NaCl, 4.18mg/ml MOPS (Sigma M1254), 2.22mg/ml KCl, 248μg/ml NH_4_Cl was adjusted to pH 6.8 using HCl or NaOH and filter sterilized. 2X agar base containing 30g/L agar, 10mg/ml porcine mucin Type II (Sigma M2378) was autoclaved to sterilize. The 2X buffered base and 2X agar base were combined and additional sterile ingredients were added to a final concentration of 3.03mM glucose (stock 1.11M), 9.3mM L-lactic acid (stock 1M), 1.75mM CaCl_2_ dihydrate (stock 1M), 0.606mM MgCl_2_ hexahydrate (stock 1M), 0.3mM N-acetylglucosamine (stock 0.25M), 0.036mM FeSO_4_ heptahydrate (stock 0.036M), 0.066mM tryptophan (stock 0.1M), and 0.136mM 1,2-dipalmitoyl-sn-glycero-3-phosphocholine (stock 0.34M in chloroform).

Quintuplicate colony biofilms were grown for 16 h at 37°C. Colonies were scraped from the agar surface using a rubber policeman, deposited in a 1.5 ml tube, stored at -80°C, and sent to Metabolon for metabolomics analysis. Values were scaled and imputed, then normalized using the total raw reads across all metabolites for a given strain, then re-scaled. A Welch’s t-test (two sample, unequal variance) was used for statistical analysis of each metabolite, and values with a p-value less than 0.1 were accepted as statistically significant. Fold-change ratios (Δ*lasR*/Δ*lasR*Δ*mhr*) were calculated and transformed into log_2_ values. Additional details and raw data are located in **Table S1**.

## Acknowledgements

The authors would like to thank Dallas Mould for her assistance with metabolomics sample collection and submission, Jeanyoung Jo, Lars Dietrich, and Elora Demers for providing editing support, and Tom Hampton for helpful discussions pertaining to statistical analysis.

