## Supplemental Data 1 for "*Pseudomonas aeruginosa lasR* mutant fitness in microoxia is supported by an Anr-regulated oxygen-binding hemerythrin"

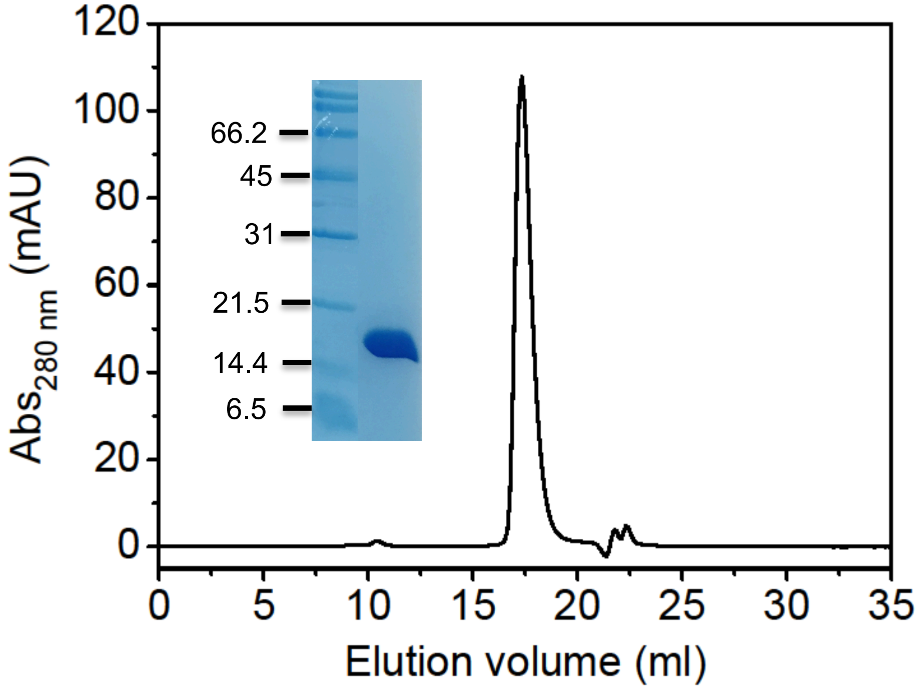

**Figure S1: Size exclusion chromatography profile and the SDS-PAGE gel (inset) of Mhr expressed in and purified from *E. coli*.** The Mhr was recovered at 17.37 ml elution volume, yielding the molecular weight of ~18.6 kDa according to the column calibration.

| | gi | | |
|---|---|---|---|
| Consensus | | ..................D.QH...................................H...EE.......Y.....H...H | |
| | | 22 ... 58 ... 62 ... 77 ... 81 | |
| Pseudomonas_aeruginosa | 9947645 | -MAHLVWQDDLNTGIQVIDNQHKRIVEMINHLHDAQQ-GKEHAAIAEVIEELVDYTLSHFAFEETLMEDAGYQFSRAHKKIH | 80 |
| Pseudomonas_citronellolis | 1004378340 | -MAYLVWQDDLNTGIQVIDNQHKRIVEMINQLHEAQG-TLDKARVGEVIEELVDYTQSHFAFEETLLEDSGYQFTRAHKKVH | 80 |
| Pseudomonas_resinovorans | 1148798354 | -MVYLEWESDLDTGIEVIDGQHKRIVEMINGLHAAQL-RMDKAAVARVIEEVVDYTLSHFAFEETLLEDAGYQFSRAHKKVH | 80 |
| Pseudomonas_mendocina | 1443027757 | -MAYLEWSDDLDTGIRVIDDQHRRIVAMINQLDDAQR-TGSKAKVATVIDELIDYTVSHFAFEEAMLEEAGYVFTKAHKRVH | 80 |
| Pseudomonas_pseudoalcaligenes | 1481196702 | -MAYLDWSDDLNTGITVIDDQHKRIVAMINQLDAAQR-AASKVQAGEVIDELIDYTVSHFAFEEAMIEEAGYIFTKAHKRVH | 80 |
| Pseudomonas_oryzae | 1222297357 | MGYVINWSDEYATGIDIIDEQHKRIFEYLSEIDHAIK-AQSLADVEHVVKAVVDYAISHNTFEESLMEKAGYPMLDAHHKVH | 81 |
| Azotobacter_chroococcum | 748252646 | -MAYLTWRNDLNTGIEVIDAQHRQIVEMINQLHETQ--GQDRAAVGVVIEALVDYTVSHFAFEESLMEDAGYQFSRAHKRIH | 79 |
| Dyella_sp | 1433330587 | -MTALVWQEDLNTGIDVIDNQHRRIVAIVNALSDAQQ-RQDRRAVGEVLEELVDYTLSHFTFEEALMEDAGYEFGRPHKRLH | 80 |
| Stenotrophomonas_maltophilia | 1407328191 | -MALLVWQDDLNIGIDVIDQQHRRIIEMLNHLHVAQA-SMQRAAVGEVIDEVVDYTMSHFAFEEELMEEAGYPFCAAHKRVH | 80 |
| Xanthomonas_campestris | 1451663768 | -MALLIWQDELNTGIEVIDHQHRRIVEMINHLYVAQT-SLQRLVVSDVIDELIDYTLSHFAFEEELMEEAGYPFSHAHQRVH | 80 |
| Acinetobacter_baumannii | 1479469231 | --MKMKWASEYNTGIDVIDEQHKRILDYINEIDDVKD-DEDRRRIKDVLDNIIDYTQSHFTFEESLQEEADYKYRVPHKRVH | 79 |
| Campylobacter_jejuni | 1444892534 | ---MTYNEKIISMNNDLLDHQHKELFEISKKLSLMNQRHVGTKELKIVLRELLIMINRHFSDEEAFMREIEYPYINHHTRIH | 79 |
| Methylococcus_capsulatus | 81682690 | MALMTWTAAEFGTNVGFADDQHKTIFDMVNKLHDTAA-TGNRSEIGKQLDALIDYVVMHFKSEETEMQKKGYADFAAHKAEH | 81 |

| | Identity percentage | | |
|---|---|---|---|
| Consensus | | .....................................H....D.................................. | |
| | | 117 ... 122 | |
| Pseudomonas_aeruginosa | 100% | ELFIRRVSEYRVRFQAGED---VGDELKGLLSRWLFNHIRNDDAGYVDAVRHSMSELVKDKSEGGWLSRSMKRFFG | 153 |
| Pseudomonas_citronellolis | 77.8% | ELFIKRVSDYRLRFAAGED---VAEELKGLLGRWLFTHIRNDDANYVESVKASMHELTHDQSHGGWLSRSMKRFFG | 153 |
| Pseudomonas_resinovorans | 72.5% | ELFIRRVNVFRTRHQAGDD---VADELKDLLGRWLFNHIRNDDANYVEAVKANMQQLTRDDSNSGWLSRSMRRFFG | 153 |
| Pseudomonas_mendocina | 61.4% | ALFIKRVEDYRQRFTSGED---IADELKGLLGRWLFSHIRSDDRNYVEAVNDNLRRLSADASEGGWLRRARRRFFRGAA | 156 |
| Pseudomonas_pseudoalcaligenes | 60.8% | ALFIKRVEDYRERFNRGED---IADELKGMLGRWLFSHIRSDDRNYVEAVNENLRRLSADASEGGWLRRAGRRFFRGAA | 156 |
| Pseudomonas_oryzae | 43.4% | ESFKARVNGYLQRLNGGEDAFRLAREVRGDLGLWLTNHIKRDDRHYVPYVKK----S-LD---GGFVSRMLGKFFR | 149 |
| Azotobacter_chroococcum | 70.4% | DLFIKRVSDYRGRYAAGED---VAEELQELLSRWLFSHIKEEDASYVGAVRAQMQSLASDRSSGGWLSRSMKRFFGRSEAA | 157 |
| Dyella_sp | 64.7 | DIFAARIRQYQLRFAEGED---VADELKSMLSRWLFQHIRNDDRAYVESVRRNMLKLTSDTSEGGWLSVALKRFFGRKGA | 157 |
| Stenotrophomonas_maltophilia | 63.4% | EVFIKRVSEYRMRFQAGED---ISDELRTMLSRWLFNHIRGDDQAYAEQVKAHLNQFAREHQGGSWLGRTLKRFFG | 153 |
| Xanthomonas_campestris | 58.8% | AVFAKRVGDYRLRFQAGED---VSDELRNMLSRWLFNHIRGDDQAYPPQVIAHLDQFSRTHQHGSWLGRTLKRLFR | 153 |
| Acinetobacter_baumannii | 41.7% | DLFIKKIELYRERFEMGHT---IEAELQEILAKWLINHIQHDDADYVGAVKENMMGIIREK-EKKKGKNWFARFFS | 151 |
| Campylobacter_jejuni | 19.8% | RKIILEIEEIISEAKFVN--IMTEKLNLVVQDFIFKHTAKEDSKIVKYYEEKFKK | 133 |
| Methylococcus_capsulatus | 30.2% | DKLVGVCADLQKKFHAGEA--EVNQDTTRFVRDWLVNHIPKVDKLYGPCLSA | 131 |

**Figure S2. Alignment of Mhr homologs from different bacteria and conservation of the di-iron binding motif.** The conserved residues, present in the hemerythrin active site (see Figure 3), are highlighted in yellow.

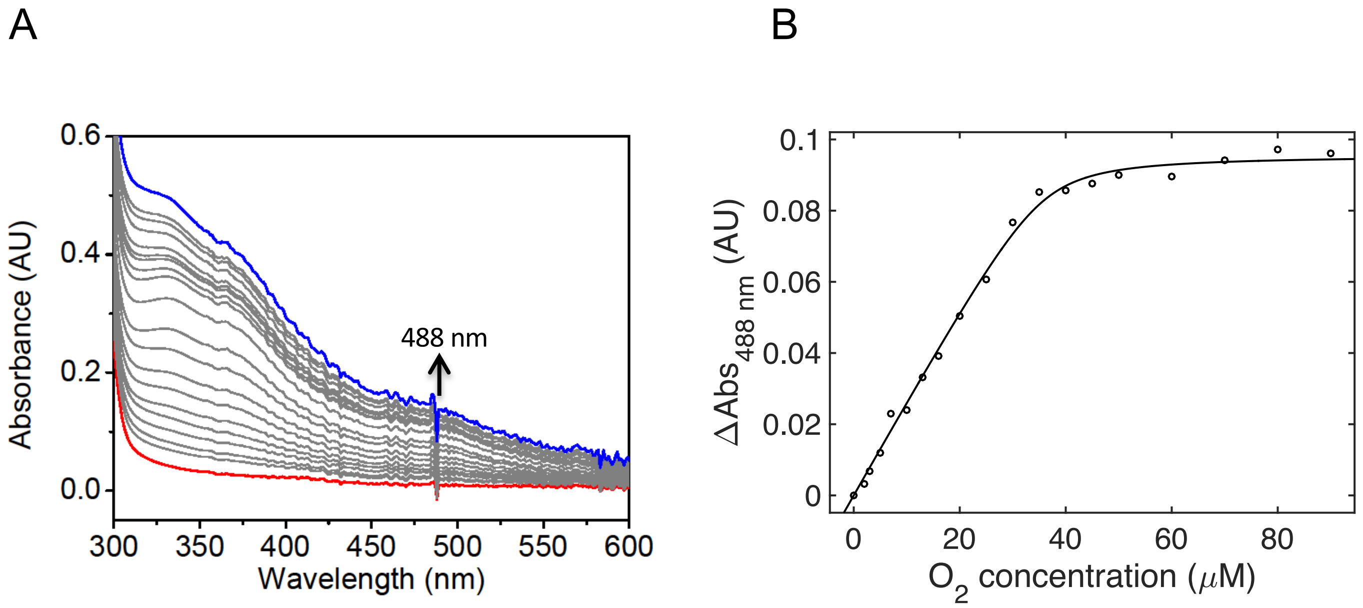

**Figure S3. Analysis of Mhr $O_2$ binding**. A) Electronic absorption spectral change during the process of $O_2$ titration into Mhr. The absorbance increase at 488 nm indicates the formation of Oxy-Mhr. B) The titration curve of the absorbance at 488 nm against the total $O_2$ concentration was fitted as described in the Materials and Methods, which yielded the value $K_d$ = 0.74 μM.

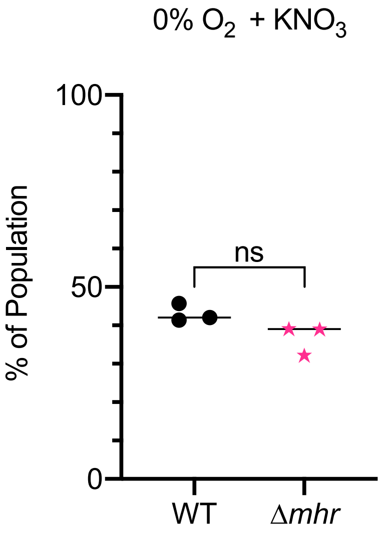

**Figure S4. Relative fitness of the Δ*mhr* mutant in anoxic denitrifying conditions.** Fitness assay using WT and Δ*mhr* on tryptone agar supplemented with 100 mM $KNO_3$ in anoxic conditions.

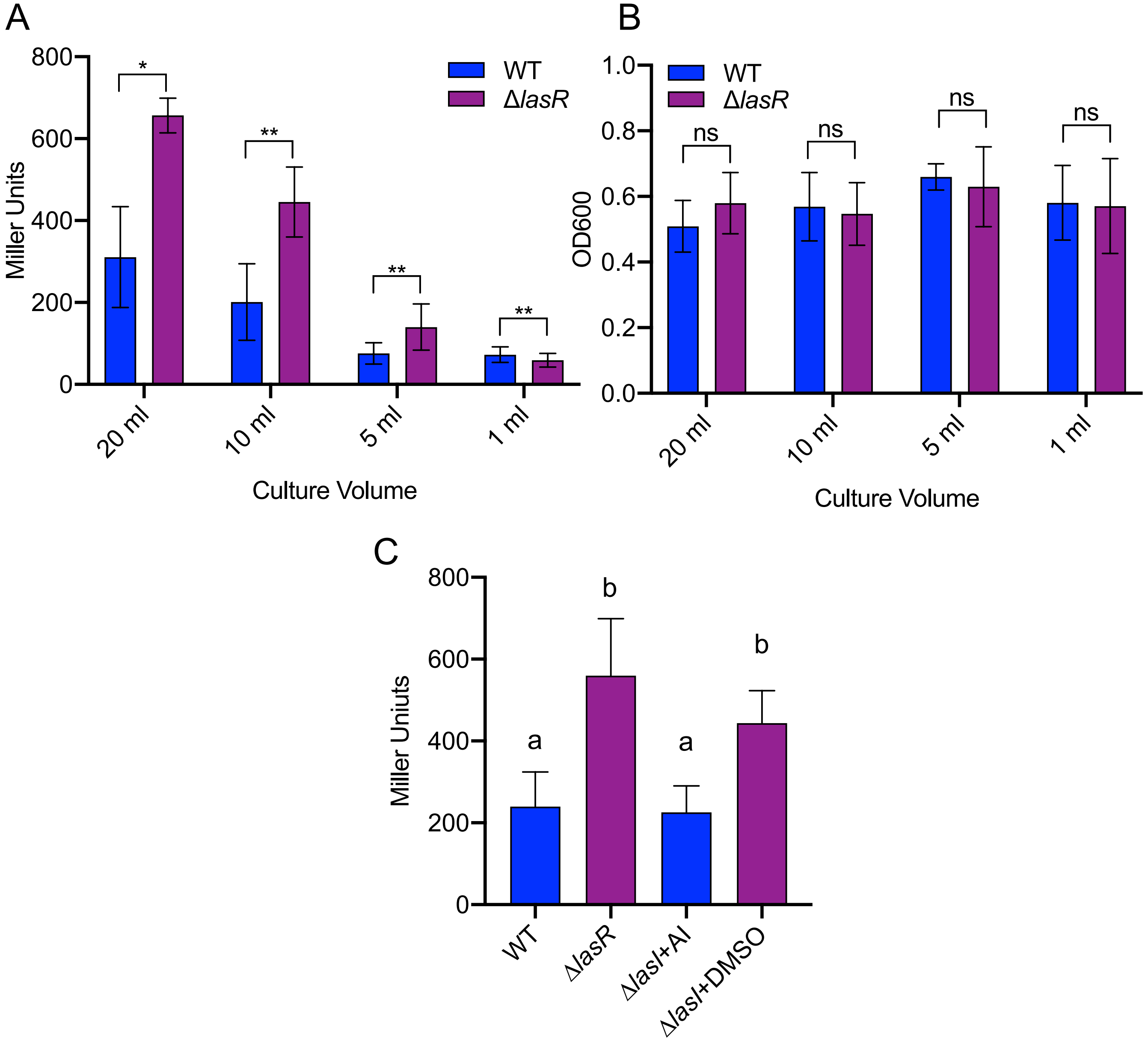

**Figure S5: Anr activity in the absence of LasR signaling.** *mhr* promoter activity was measured in cells from cultures grown in 50 ml capacity flasks containing different volumes of medium (1, 5, 10, 20 ml) with shaking (250 rpm) for 6 h at 37°C. A) *mhr* promoter activity in wild-type *P. aeruginosa* PA14 (WT) and the Δ*lasR* mutant. B) Final culture density of flasks in A) as measured by $OD_{600}$. Significance of the differences between strains was compared using a ratio paired t-test for each volume condition. C) The *mhr* promoter activity was measured by ONPG assay in WT, Δ*lasR*, and Δ*lasI* mutant with either 10 μM 3OC12HSL autoinducer (AI) or equivalent volume of vehicle (DMSO). Ten ml culture volumes were used. For A and B, bars represent averages across at least three different experiments performed on different days. (*p<0.05, **p<0.01 by ratio paired t-test for each volume condition). For C, data represent averages of four independent experiments. Samples with different letters are statistically different by two-way ANOVA, p<0.05.

**Table S2: Strains, plasmids, and primers used in this study.**

| Strain | Strain ID | Description | Source |
|---|---|---|---|
| ***P. aeruginosa*** | | | |
| PA14 WT p*mhr-lacZ* | DH2857 | DH122 with reporter for activity at the *mhr* promoter (GH121 p*mhr-lacZ*) | This study |
| PA14 Δ*lasR* p*mhr-lacZ* | DH3544 | DH164 with reporter for activity at the *mhr* promoter (GH121 p*mhr-lacZ*) | This study |
| PA14 Δ*anr* p*mhr-lacZ* | DH2936 | DH2855 with reporter for activity at the *mhr* promoter (GH121 p*mhr-lacZ*) | This study |
| PA14 WT p*mhr(*Mut.)-*lacZ* | DH2884 | DH122 WITH Reporter for activity at the *mhr* promoter, Anr box mutated (GH121 p*mhr* (Mut.)-*lacZ*) | This study |
| PA14 WT | DH122 | Wild type *P. aeruginosa* | [1] |
| PA14 Δ*lasR* | DH164 | DH122 with in frame deletion of *lasR* (PA14_45960) | [2] |
| PA14 WT att::*lacZ* | DH22 | PA14 WT with constitutive expression of *lacZ* | Roberto Kolter |
| PA14 WT att::*lacZ* + pMQ70 | DH3545 | DH3545 carrying pMQ70 | This study |
| PA14 Δ*mhr* | DH2853 | DH122 with in frame deletion of *mhr* (PA14_42860) | This study |
| PA14 Δ*mhr* + EV | DH2861 | DH2853 carrying empty vector for gene expression (pMQ70) | This study |
| PA14 Δ*mhr* + pMQ70_*mhr* | DH2863 | DH2853 with inducible expression of *mhr* (pMQ70_*mhr*) | This study |
| PA14 Δ*lasR*Δ*mhr* | DH2917 | DH164 with in frame deletion of *mhr* (PA14_42860) | This study |
| PA14 Δ*lasR*Δ*mhr* + EV | DH3546 | DH2917 carrying empty vector for gene expression (pMQ70) | This study |
| PA14 Δ*lasR*Δ*mhr* + pMQ70_*mhr* | DH3547 | DH2917 with inducible expression of *mhr* (pMQ70_*mhr*) | This study |
| PA14 Δ*cco* | DH2865 | DH122 with in frame deletion of *ccoNOQP-1*, *ccoNOQP-2*, and all DNA between the two operons (PA14_44340-PA14_44400) | This study |
| PA14 Δ*mhr*Δ*cco* | DH2866 | DH2853 with in frame deletion of *ccoNOQP-1*, *ccoNOQP-2*, and all DNA between the two operons (PA14_44340-PA14_44400) | This study |
| PA14 Δ*cco* att::lacZ | DH2911 | DH22 with in frame deletion of *ccoNOQP-1*, *ccoNOQP-2*, and all DNA between the two operons (PA14_44340-PA14_44400) | This study |
| PA14 ΔlasR + lasR | DH3549 | DH164 with complementation of *lasR* (PA14_45960) at the native locus | This study |
| PA14 Δanr | DH2855 | DH122 with in frame deletion of *anr* (PA14_44490) | This study |
| PA14 Δanr + anr | DH3478 | DH2855 with complementation of *anr* (PA14_44490) at the native locus | This study |
| PA14 ΔlasRΔanr | DH2401 | In frame deletions of *lasR* (PA14_45960) and *anr* (PA14_44490) | [3] |
| PA14 ΔlasRΔanr + anr | DH3479 | DH2401 with complementation of *anr* at the native locus | This study |
| 262K | DH2590 | Acute keratitis clinical isolate; *lasR*$^{I215S}$ | [4] |

| | | | |
|---|---|---|---|
| 262K + *lasR*$^{PA14}$ | DH2743 | DH2415 with PA14 *lasR* allelic replacement at the native locus | This study |
| NC-AMT0101-1-2 | DH2417 | Chronic lung infection isolate with functional LasR allele, parent of NC-AMT0101-1 | [5] |
| NC-AMT0101-1-1 | DH2415 | Chronic lung infection isolate related to DH2417 with LasR LOF (frame shift) allele | [5] |

***E. coli***

| | | |
|---|---|---|
| S17 λ | | Used as a conjugation partner for introducing pMQ30-based plasmids. |

| **Plasmid** | **Strain ID** | **Description** | **Source** |
|---|---|---|---|
| GH121 | DH2830 | For inserting sequences at the att::Tn7 site via allelic replacement; $Gm^R$ | [6] |
| GH121 p*mhr-lacZ* | DH2856 | lacZ under control of the mhr promoter, integrated at the att::Tn7 site; $Gm^R$ | This study |
| GH121 p*mhr* (Mut.)-*lacZ* | DH2876 | lacZ under control of the mhr promoter with a mutated Anr box, integrated at the att::Tn7 site; $Gm^R$ | This study |
| pMQ30 | DH962 | Shuttle vector for yeast cloning and Gram-negative allelic replacement; $Gm^R$ | [7] |
| pMQ30 Δ*cco* | DH2919 | For deleting *ccoN1O1Q1P1*, *ccoN2O2Q2P2*, and all DNA between the two operons (PA14_44340-PA14_44400) | This study |
| pMQ70 | DH2932 | Shuttle vector for yeast cloning and arabinose-inducible gene expression; $Amp^R$ | [7] |
| pMQ70_*mhr* | DH2858 | Vector for arabinose-inducible gene expression of *mhr*; $Amp^R$ | This study |
| pMQ30 Δ*mhr* | DH2854 | For deleting *mhr*; $Gm^R$ | This study |
| *pMQ30 +anr* | DH3481 | For complementing *anr* at the native locus; $Gm^R$ | [3] |
| pMQ30 +*lasR* | DH3548 | For complementing *lasR* at the native locus; $Gm^R$ | This study |
| pET20b(+)_Pa *mhr* | DH3598 | For protein expression and purification of Mhr | This study |

| Plasmid Constructed | Primer Sequence | |
|---|---|---|
| pMQ70_*mhr* | 5’-GGGAATTCCATATGGCGCATCTGGTCTGGCAGG-3’ | This study |
| pMQ70_*mhr* | 5’-CCCGCTCGAGCGACTGAAAATACAAATTCTCGCCGAAGAAGCGCTTCATCGAGCG-3’ | This study |
| pET20b(+)_*mhr* | 5’-GGGAATTCCATATGGCGCATCTGGTCTGGCAGG-3’ | This study |
| pET20b(+)_*mhr* | 5’-CCCGCTCGAGCGACTGAAAATACAAATTCTCGCCGAAGAAGCGCTTCATCGAGCG-3’ | This study |
| GH121 p*mhr-lacZ* | 5-'GCGGGCGTCGCGATCGCCGGGGCCGCATGACTGCCCACCACTGGTGACGCTGCTTTCCG-3' | This study |
| GH121 p*mhr-lacZ* | 5'-CCAGTGAAAAGTTCTTCTCCTTTACTCATGATGACTTCCCCAAGCGAACAGCG-3' | This study |
| GH121 p*mhr-lacZ* | 5'-CGCTGTTCGCTTGGGGAAGTCATCATGAGTAAAGGAGAAGAACTTTTCACTGG-3' | This study |
| GH121 p*mhr-lacZ* | 5'-CCGTCGAGTAACGCCGGCCGCGTACCCGAGAAGCCGGGTTATTATTATTTTTGACACCAGACCAACTGG-3' | This study |
| GH121 p*mhr* (Mut.)-*lacZ* | 5'-TTGTGACGCAGTACGGGTACTGACGCCGGTCAGGCGGGACAAGATGTTAATGAATTTCGC-3' | This study |
| GH121 p*mhr* (Mut.)-*lacZ* | 5'-ATCTTGTCCCGCCTGACCGGCGTCAGTACCCGTACTGCGTCACAAAAATACCCTGGCCCC-3' | This study |
| pMQ30 Δ*cco* | 5'-GACCGCTTCTGCGTTCTGATTTAATCTGTATCAGGCTGAGAACCGCCGAGAAAGCCGGTG-3' | This study |
| pMQ30 Δ*cco* | 5'-GTCGTGGGGGAGGGGAGCGCGCTATGGGCTTCCATCCACGGTTATAC-3' | This study |
| pMQ30 Δ*cco* | 5'-GTATAACCGTGGATGGAAGCCCATAGCGCGCTCCCCTCCCCCACGAC-3' | This study |
| pMQ30 Δ*cco* | 5'-TTGTGAGCGGATAACAATTTCACACAGGAAACAGCTATGTCGAGGGACTTCTTGCGCGGG-3' | This study |
| pMQ30 Δ*mhr* | 5'-TAACGCCAGGGTTTTCCCAGTCACGACGTTGTAAAACGAAAGCAGCTCAGCCGGGGCCG-3' | This study |
| pMQ30 Δ*mhr* | 5'-TCAGCCGAAGAAGCGCTTCATCGAGCGCGCAGGTCATCCTGCCAGACCAGATGCGCCAT-3' | This study |
| pMQ30 Δ*mhr* | 5'-ATGGCGCATCTGGTCTGGCAGGATGACCTGCGCGCTCGATGAAGCGCTTCTTCGGCTGA-3' | This study |
| pMQ30 Δ*mhr* | 5'-CGAGCTCGGTACCCGGGGATCCTCTAGAGTCGACCTGCAGCAAGCTCAAGCAGGGCCTG-3' | This study |
| pMQ30 +*lasR* | 5'-CAGACCGCTTCTGCGTTCTGATTTAATCTGTATCAGGCTGACCTGCCGATGACGCCGGCG-3' | This study |
| pMQ30 +*lasR* | 5'-CTCAAGAAAACCGTCAACCAAGGCCATAGCGCTACGTTCTTCTTAAACTATTAACCAATC-3' | This study |
| pMQ30 +*lasR* | 5'-ATTGGTTAATAGTTTAAGAAGAACGTAGCGCTATGGCCTTGGTTGACGGTTTTCTTGAGC-3' | This study |
| pMQ30 +*lasR* | 5'-TCGCCAGCTCGCCGACCTGAGAGGCAAGATCAGAGAGTAATAAGACCCAAATTAACGGCC-3' | This study |
| pMQ30 +*lasR* | 5'-CCATTATGGCCGTTAATTTGGGTCTTATTACTCTCTGATCTTGCCTCTCAGGTCGGCGAG-3' | This study |
| pMQ30 +*lasR* | 5'-TGAGCGGATAACAATTTCACACAGGAAACAGCTATGCGTAAAGCGCGATCTGGGTCTTGG-3' | This study |
